# Manure substitution of chemical fertilizer affect soil microbial community diversity, structure and function in greenhouse vegetable production systems

**DOI:** 10.1101/570986

**Authors:** Haoan Luan, Wei Gao, Shaowen Huang, Jiwei Tang, Mingyue Li, Huaizhi Zhang, Xinping Chen

## Abstract

Soil microbial community and enzyme activities together affect various ecosystem functions of soils. Fertilization, as important agricultural management practices, are known to modify soil microbial characteristics; however, inconsistent results have been reported. The aim of this research therefore was to make a comparative study of the effects of different fertilization patterns (No N inputs (No N), 100% chemical fertilizer-N (CN) inputs (4/4CN) and different substitution rates of CN by organic manure-N (MN) (3/4CN+1/4MN, 2/4CN+2/4MN and 1/4CN+3/4MN)) on soil physicochemical properties, enzyme activities and microbial attributes in a GVP of Tianjin, China. Manure substitution of chemical fertilizer, especially at higher substitution rate (2/4CN+2/4MN and 1/4CN+3/4MN), improved soil physicochemical properties (higher soil organic C (SOC) and nutrient contents; lower bulk densities), promoted microbial growth (higher total phospholipid fatty acids and microbial biomass C contents) and activity (higher soil hydrolase activities). Manure addition caused a remarkable increase of the fungi/bacteria ratio and a distinct shift in the fungal (bacterial) community to greater abundance of arbuscular mycorrhizal fungi (G+ bacteria) compared with saprotrophic fungi (G− bacteria). These changes drove shifts toward fungal-dominated soil microbial communities and then optimized microbial community structure. Also, manure application increased soil biodiversity (microbial community and enzyme function), indicated by increased Shannon–Wiener diversity. Redundancy analysis indicated that the most possible mechanism of the impacts of different fertilization patterns on soil microbial characteristics may be the mediation of SOC and nutrient (N) availability (especially SOC) in this GVP of China. In conclusion, manure substitution of chemical fertilizer, especially at higher substitution rate, was more efficient for improving soil quality and biological functions.

## 1. Introduction

Vegetable production is a vital part of agricultural sector in China. In 2015, the areas of vegetable planting in China were 2.1 × 10^7^ ha [1], which was around 9.5 and 210.0 times, respectively, of that in Europe (2.2 × 10^6^ ha [2]) and Canada (1.0 × 10^5^ ha [3]). Over the last 30 years, greenhouse vegetable productions (GVPs) in China have grown rapidly and become the main type of vegetable production due to its higher economic benefits relative to vegetable productions in open-air fields [4]. As a high-intensity agricultural ecosystem, the GVPs has been characterized as large cropping index, high agricultural inputs (e.g., pesticides, chemical and organic fertilizers), closed or semi-closed environment with higher inside temperature and humidity, which has led to a series of soil ecological and biological problems [5, 6].

Soil microorganisms are important parts of agricultural ecosystems and mediate many processes, including soil aggregate formation [7], soil organic C (SOC) decomposition and nutrient transformation [8] in agricultural systems. Therefore, they further regulate soil fertility and agricultural productivity [9]. Soil extracellular enzymes, produced and secreted by soil microorganisms, provide a functional fingerprint of soil microbial community because they are often involved in SOC formation and nutrient cycling [10]. Previous studies have demonstrated that soil microorganisms and extracellular enzymes are related to many soil properties, such as soil moisture [11], temperature [12], SOC [13], C/N ratio [14], nutrient status [15] and pH [16], and which factors become dominant depends on differences in climate and soil type.

In agricultural production systems, different fertilization regimes (manure vs. chemical fertilizer) have been found to affect soil physical, chemical and biological properties in different ways [17]. Chemical fertilizers, especially nitrogen (N) inputs, have been major contributors to the impressive crop yield increases realized since the 1950s [18]. However, excessive chemical fertilization leads to soil quality degradation and environmental pollution in recent years [19]. For solving these problems, organic amendments and substitution of chemical fertilizers are increasingly recommended as effective and sustainable practices to sustain crop yield and soil quality [20, 21]. Manure has been found to be effective to increase microbial biomass and activities in soil, and modifies soil microbial community [22]. Nevertheless, the effects of manure application on soil microorganisms varied with soil type, quantity and quality of manure and other factors [23, 24]. To our knowledge, there is no comprehensive study exploring the changes of soil microbial properties under different fertilization regimes in GVPs. Thus, the impact of manure application on soil microbial properties and the underlying mechanisms still need to be investigated in GVPs.

The objectives of this study were (1) to investigate the effects of 8 years of continuous manure application on soil physicochemical properties, soil microbial community composition and soil extracellular enzyme activities, and (2) to elucidate the main controlling environmental factors that drive the changes in soil enzyme activities or microbial community compositions in GVP systems. We hypothesized that manure application would improve soil physicochemical properties and increase soil microbial biomass and activity, and that these positive effects of manure application would be enhanced with manure inputs.

## 2. Materials and methods

### 2.1 Study site

A field experiment with celery (Apium graveolens cv. Wentula) -tomato (Lycopersicon esculentum cv. Chaoyan No. 299) rotation system was started in October 2009 at a solar greenhouse farm in Xiqing District, Tianjin City, China (117°0’E, 39°13’N). This region has a warm sub-humid continental climate. The mean annual temperature is 11.6°C with mean annual precipitation of 586 mm. The frost-free period is 203 days and the annual sunshine duration is 2810 h. The soils in the experimental site are medium-loam Chao (Aquic cambisols) soil (FAO classification) with the groundwater depth of 100 cm. Some of the initial physicochemical characteristics of the surface (0-20 cm) soil were as follows: pH 7.9, SOC 14.7 g kg^−1^, NO_3_^−^-N 86.2 mg kg^−1^, available P 144.6 mg kg^−1^, and available K 404.0 mg kg^−1^.

### 2.2. Experimental design

This study comprised five treatments as follows: (1) no chemical N (CN) inputs (No N), (2) 100% CN inputs (4/4CN), (3) 75% CN plus 25% manure N (MN) inputs (3/4CN+1/4MN), (4) 50% CN plus 50% MN inputs (2/4CN+2/4MN), and (5) 25% CN plus 75% MN inputs (1/4CN+3/4MN). The same amounts of N, P_2_O_5_, and K_2_O (N, P_2_O_5_ and K_2_O were 450.0, 225.0 and 600.0 kg ha^−1^ in the tomato season, respectively, and 450.0, 300.0 and 600.0 kg ha^−1^ in the celery season, respectively) were added to each treatment except the No N treatment. A detailed description of the N and C inputs is shown in Table 1. The chemical fertilizers were urea (N 46%), calcium superphosphate (P_2_O_5_ 12%), diammonium phosphate (N 18%, P_2_O_5_ 46%), potassium chloride (K_2_O 60%) and monopotassium phosphate (P_2_O_5_ 52%, K_2_O 34%). The commercial pig manure (i.e., organic manure) had a C content of 218.0 g kg^−1^, N content of 21.7 g kg^−1^, P_2_O_5_ content of 13.9 g kg^−1^, K content of 16.3 g kg^−1^ (by dry weight) and its water content was 28.9%.

Manure was applied as basal fertilization. During the tomato season, chemical fertilizers (20% N, 70% P_2_O_5_, and 20% K_2_O) were applied as basal fertilization. The remaining N and K_2_O were applied as top dressings in four doses (30% N and 10% K_2_O in the flowering period, then 30% N and 30% K_2_O, 10% N and 30% K_2_O, and 10% N and 10% K_2_O in the first, second, and third fruit cluster expansion periods, respectively). The remaining P_2_O_5_ was applied as top dressing in two doses (15% P_2_O_5_ in the flowering period and 15% P_2_O_5_ in the first fruit cluster expansion period). During the celery season, the same amount of chemical fertilizer as tomato season was applied as basal fertilization. The remaining N and K_2_O were applied as top dressings in three doses (35% N and 10% K_2_O, 35% N and 35% K_2_O, and 10% N and 35% K_2_O in the 5–6 leaf, 8–9 leaf, and 11–12 leaf periods, respectively). The remaining P_2_O_5_ (30%) was applied as a top dressing in the 5–6 leaf period. The basal fertilizer was broadcast and then tilled into the soil using a rotary tilling device, and the top dressing fertilizers were dissolved in water and then poured onto the soil.

A randomized block design was used, and each treatment was performed in triplicate. Each plot was 14.4 m^2^ (2.4 m × 6.0 m), and neighboring plots were separated with PVC plates 105 cm deep (100 cm underground, 5 cm aboveground) to avoid nutrients and water being transferred between neighboring plots. The tomato planting density was 25000 plants ha^−1^, with rows 0.3 m apart and plants 0.6 m apart. The celery planting density was 330570 plants ha^−1^, with rows 0.15 m apart and plants 0.2 m apart. Each plot was fitted with a water meter to ensure the amount of irrigation applied was accurately controlled. If the field capacity dropped below 60%, we irrigated the soil to replenishing water to 75% of field capacity. The total irrigation amounts during the celery and tomato growing periods were 3334 and 3889 m^3^ ha^−1^ year^−1^.

### 2.3 Soil sampling and analysis

After the celery (the fifteenth-season) harvest in January 2017, ten soil cores (3 cm in diameter, 0–20 cm depth) were collected in each plot and mixed to create one combined soil sample. The fresh soil samples were sieved through a 2-mm mesh and divided into two subsamples. One subsample was air dried and then passed through 0.25-mm mesh for the determination of soil physicochemical properties, the other subsample was stored at 4°C for the measurement of soil extracellular enzyme activities, soil microbial biomass and community composition.

### 2.4 Soil physicochemical analysis

Soil pH was determined with a compound electrode (PE 10, Sartorius, Goettingen, Germany) using a soil/water ratio of 1:2.5(w/v; g cm^−3^). Electrical conductivity (EC) was measured at 25°C in 1:5 soil-water mixtures. Soil bulk density (BD) was measured after drying the soil cores at 105°C for 48 h. Soil organic carbon (SOC) and total nitrogen (TN) were determined by the heated dichromate/titration method [25] and the Semimicro-Kjeldahl method [26], respectively. Soil nitrate-N (NO_3_^−^-N) and ammonium-N (NH_4_^+^-N) was extracted by 2 M potassium chloride and measured using a flow injection autoanalyzer (Smartchem 200, Alliance, France). Soil available phosphorus (AP) was extracted by 0.5 M sodium bicarbonate (pH 8.5) and determined by the Olsen method [27]. Soil available potassium (AK) was extracted by 1 M ammonium acetate, adjusted to pH 7.0, and then measured by atomic absorption spectrometry (NovAA300, Analytik Jena AG). Soil microbial biomass carbon/nitrogen (MBC/MBN) were measured by the chloroform fumigation-extraction method [28]. Moreover, the values of MBC and SOC were used to calculate the microbial quotient (MQ, the ratio of MBC to SOC), which could be used as indicators of soil microbial activity [29].

### 2.5 Phospholipid fatty acid analysis

Soil microbial community composition was determined by phospholipid fatty acid (PLFA) analysis according to the procedure described by Wu et al. [30]. Briefly, the soil samples were freeze-dried, then PLFAs were extracted using a 1:2:0.8 (by volume) chloroform–methanol–citrate buffer mixture at pH 4.0. The chloroform extract was passed through a SPE-Si column (Supelco, Poole, UK), then the neutral lipids, glycolipids, and polar lipids were eluted with chloroform, acetone, and methanol, respectively. Nonadecanoic acid methyl ester (19:0) was added as an internal standard, and the recovered polar lipids were converted into fatty acid methyl esters (FAMES) by a mild alkaline methanolysis. Dried FAMES were redissolved in n-hexane and then quantified by gas chromatography (N6890, Agilent Technologies, Santa Clara, CA, USA) and identified with a MIDI Sherlock microbial identification system version 4.5 (MIDI Inc., Newark, DE, USA). The total and individual PLFA abundances were expressed in units of nmol g^−1^ soil. The PLFAs were divided into various taxonomic groups based on previously published PLFA biomarker data [31, 32]. Specifically, 15:00, i15:0, a15:0, i16:0, 16:1ω7c, 17:00, a17:0, i17:0, cy17:0, 18:1ω7c and cy19:0 were used to represent bacterial biomarkers [33]. The PLFA i15:0, a15:0, i16:0, a17:0 and i17:0 were used as biomarkers for Gram-positive (G+) bacteria, and the PLFA 16:1ω7c, cy17:0, 18:1ω7c and cy19:0 were designated for Gram-negative (G−) bacteria [34]. The general PLFAs (15:00 and 17:00) were summed as indicators of other bacteria [32]. The PLFAs 18:2ω6c and 18:1ω9c were attributed to saprotrophic fungi (SF), whereas PLFAs 16:1ω5c were regard as arbuscular mycorrhizal fungi (AMF) [35]. The PLFAs 10Me-16:0, 10Me-17:0 and 10Me-18:0 were calculated as indicators of actinomycete [8]. Other PLFAs such as 14:00, 16:00 and 18:00 were also included to represent the composition of microbial community [32]. Taken together, the total PLFAs of soil microbial community included all of the PLFAs mentioned above. Ratios of group-specific lipids (i.e., F/B, G+/G− and AMF/SF) were taken to reflect the relative biomass of their respective groups [36]. The ratios of cyclopropyl fatty acids to their precursors (cy17:0/16:1ω7c) and (i17:0 + i15:0) to (a17:0 + a15:0) as two bacterial physiological stress indices were evaluated in soil microbial community [37]. Moreover, soil microbial community diversity was evaluated using Shannon–Wiener diversity index (H’_M_), Pielou evenness index (J), and Margalef richness index (SR) based on the following equation [38]:

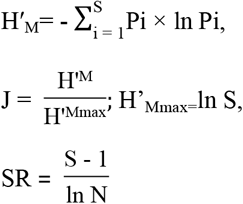

where ‘Pi’ is the percentage of the peak area of PLFA to the total area of each sample; ‘S’ is the total number of microbial PLFAs; and ‘N’ is the amount of microbial PLFAs.

### 2.6 Soil enzyme activity analysis

Seven enzymes (Table 2) involved in C and N cycling were determined using a micro-plate enzyme assay, according to the procedures of Bell et al. [39]. Briefly, 1.0 g dry-mass-equivalent of fresh soil was homogenized in 100 mL of 50 mM acetate buffer (pH 8.5). For hydrolase analysis, buffer, sample suspension, 10 mM references and 200 μM substrates (4-methylumbelliferone or 7-amino-4-methylocumarin) were dispensed into the wells of a black 96-well microplate. The microplates were covered and incubated at 25°C for 4 h in the dark, then fluorescence was measured using a microplate fluorometer (Scientific Fluoroskan Ascent FL, Thermo) with 365 nm excitation and 450 nm emission filters. Phenol oxidase and peroxidase were measured in a clear 96-well microplate using the substrate of L-3, 4-dihydroxyphenylalanine (L-DOPA). The dispensed volume and the order of buffer, sample suspension, 25 mM L-DOPA, and 0.3% (w/v) H_2_O_2_ were the same as the fluorometric enzymes. The microplates were covered and incubated at 20°C for 20 h in the dark, then the activities were determined by measuring the absorbance at 450 nm using the microplate fluorometer. The enzyme activities were expressed in nmol h^−1^ g^−1^. For each sample, the geometric mean of the assayed enzyme activities (Gmea), hydrolase (GH) and oxidase (GOR) were calculated as:

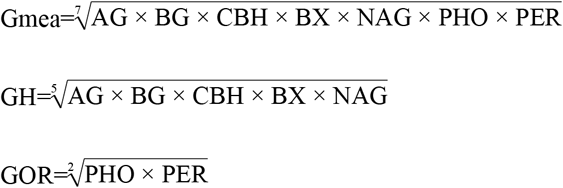

Moreover, soil enzyme functional diversity was calculated from the activities of seven enzymes using Shannon’s diversity index as [40]:

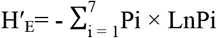

where Pi is the ratio of each enzyme activity to the sum of all enzyme activities.

### 2.7 Data analysis

The characteristics of soil samples from the plots given different fertilization treatments were analyzed using SPSS 16.0 software (SPSS Inc. Chicago, IL, USA). One-way ANOVA with Duncan tests were performed to test the significance (*P* < 0.05) of measured variables. Differences in soil enzyme activities or microbial community composition were investigated by principal component analysis (PCA) using CANOCO 4.5 (CANOCO, Microcomputer Power Inc., Ithaca, NY, USA). Redundancy analysis (RDA) was conducted to illuminate the relationships between soil microbial community, enzyme activities and physicochemical properties in different fertilization treatments using CANOCO 4.5. Monte Carlo permutation tests were performed to assess whether soil microbial community composition or enzyme activities were correlated with soil physicochemical properties.

## 3. Results

### 3.1 Changes in soil physicochemical properties

Soil pH ranged from 8.31 to 8.54 under different fertilization treatments, and soil pH under organic manure application (3/4CN+1/4MN, 2/4CN+2/4MN and 1/4CN+3/4MN) was significantly increased by 0.19-0.23 units than that under NPK fertilizer application (4/4CN) (Table 3). Soil EC was highest in 4/4CN treatment (0.49 mS cm^−1^), followed by No N treatment (0.44 mS cm^−1^) and lowest in manure-amended treatments (0.39-0.41 mS cm^−1^). Higher rates of manure input (1/4CN+3/4MN and 2/4CN+2/4MN) significantly (*P* < 0.05) decreased Soil BD by 6.40%-8.12% compared with 4/4CN treatment. The SOC, NO_3_^−^-N and NH_4_^+^-N contents were lowest in the No N treatment, while these parameters slightly increased under 3/4CN+1/MN treatment, significantly (*P* < 0.05) increased under 2/4CN+2/4MN and 1/4CN+3/MN treatments by 32.88%-50.81%, 29.94%-40.75% and 16.00%-16.79%, respectively, compared to 4/4CN treatment. The available P (AP) contents were significantly increased in manure-amended soils by 7.07%-39.88% compared with those in 4/4CN-amended soils. In contrast, soil available K (AK) content was highest in the No N treatment (542.30 mg kg^−1^) and decreased by 7.84%-17.15% upon manure addition. Moreover, with increasing manure application rates, the EC, SOC, NO_3_^−^-N, NH_4_^+^-N and AP contents increased, and the BD and AK contents decreased.

Manure-amended treatments significantly (*P* < 0.05) increased both soil MBC by 26.78%–82.91% and MBN by 73.94%–151.45% compared with chemical fertilization treatments (No N and 4/4CN) (Fig. 1a and 1b). Moreover, these parameters increased as the manure application rate increased. The MBC/MBN ratios were highest in the 4/4CN treatment (10.12), followed by the No N treatment (9.51), and the lowest values were observed in manure-amended treatments (6.94-7.38), while there were no significant differences among manure-amended treatments (Fig. 1c). The microbial quotient (MQ) values in manure-amended soils (1.83-1.96) was significantly (*P* < 0.05) higher than that in No N treatment (1.56) and slightly (*P* > 0.05) higher than that in 4/4CN treatment (1.77).

**Fig. 1.**
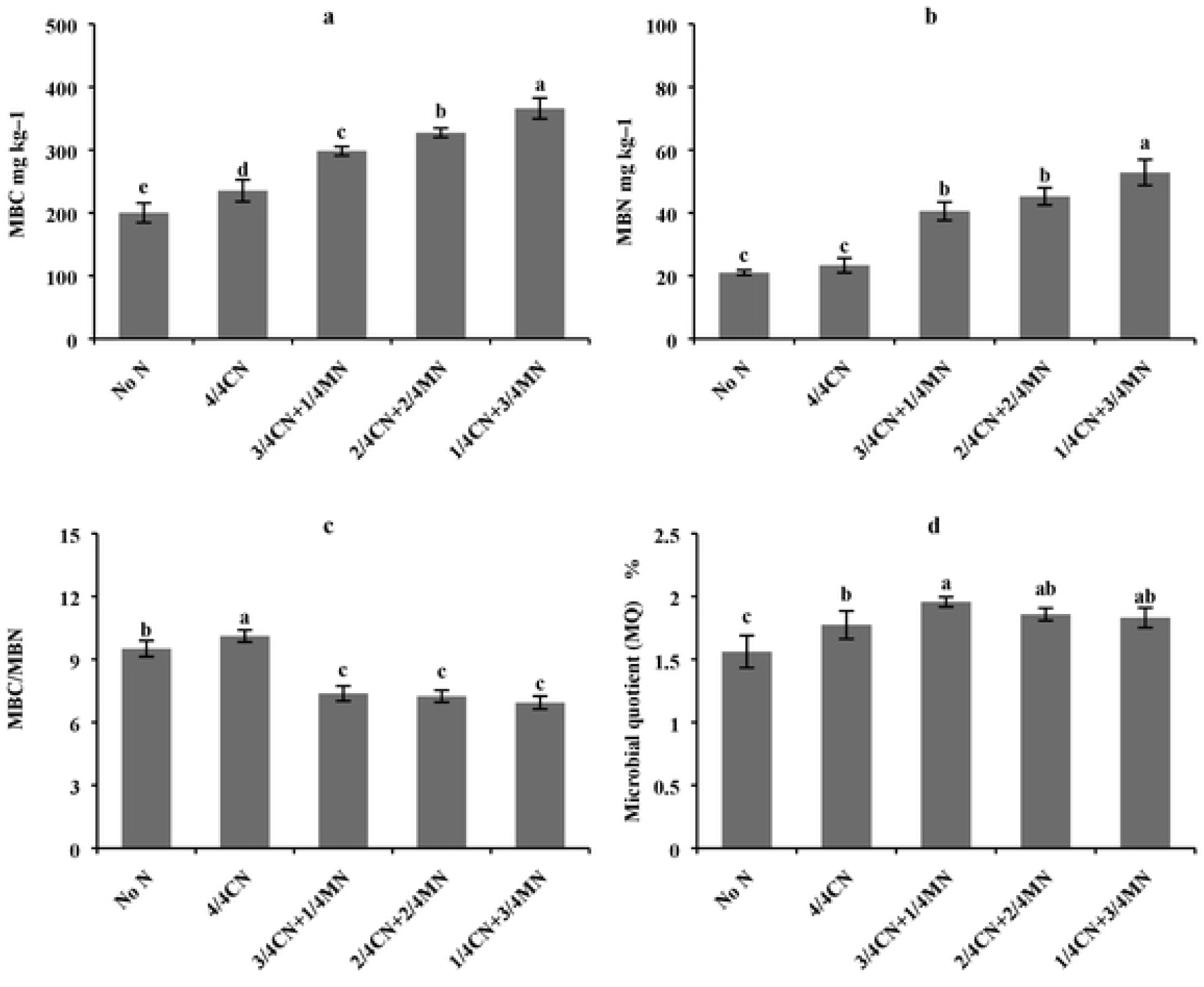
Changes in microbial biomass carbon (MBC) (a), microbial biomass nitrogen (MBN) (b), the ratio of MBC to MBN (c) and microbial quotient (MQ) (d) under different fertilization treatments after celery (the fifteenth-season) harvest in January 2017. Vertical error bars represent the standard deviation (n = 3), and lowercase letters indicate significant differences among different treatments at the *p* < 0.05 level.

### 3.2 Changes in soil enzyme activities and geometric mean of enzyme activities

Manure substitution of chemical fertilizer significantly (*P* < 0.05) affected soil extracellular enzyme activities (EEAs) with the exception of PER in this research (Fig. 2). Compared to chemical fertilization treatments, soil hydrolase (BG, CBH, NAG, BX and AG) activities were slightly higher under 3/4CN+1/MN treatment, and significantly (*P* < 0.05) higher under 2/4CN+2/4MN and 1/4CN+3/MN treatments by 23.36%-81.14%, 82.34%-196.40%, 69.95%-295.99%, 31.34%-65.69% and 23.29%-49.46%, respectively. Moreover, no significant differences were observed in soil hydrolase activities between chemical fertilization treatments. Soil oxidase activities (PHO and PER) showed different trends compared with soil hydrolase activities. The PHO activity did not differ among manure-amended treatments (2.65-2.77 nmol h^−1^ g^−1^) and No N treatment (2.77 nmol h^−1^ g^−1^), but the PHO activity was significantly (*P* < 0.05) higher in the manure-amended soils (by 6.99%–11.79%) than that in 4/4CN treated soils (2.48 nmol h^−1^ g^−1^). Additionally, there were no significant differences in the PER activity among different fertilization treatments (2.65–2.74 nmol h^−1^ g^−1^).

**Fig. 2.**
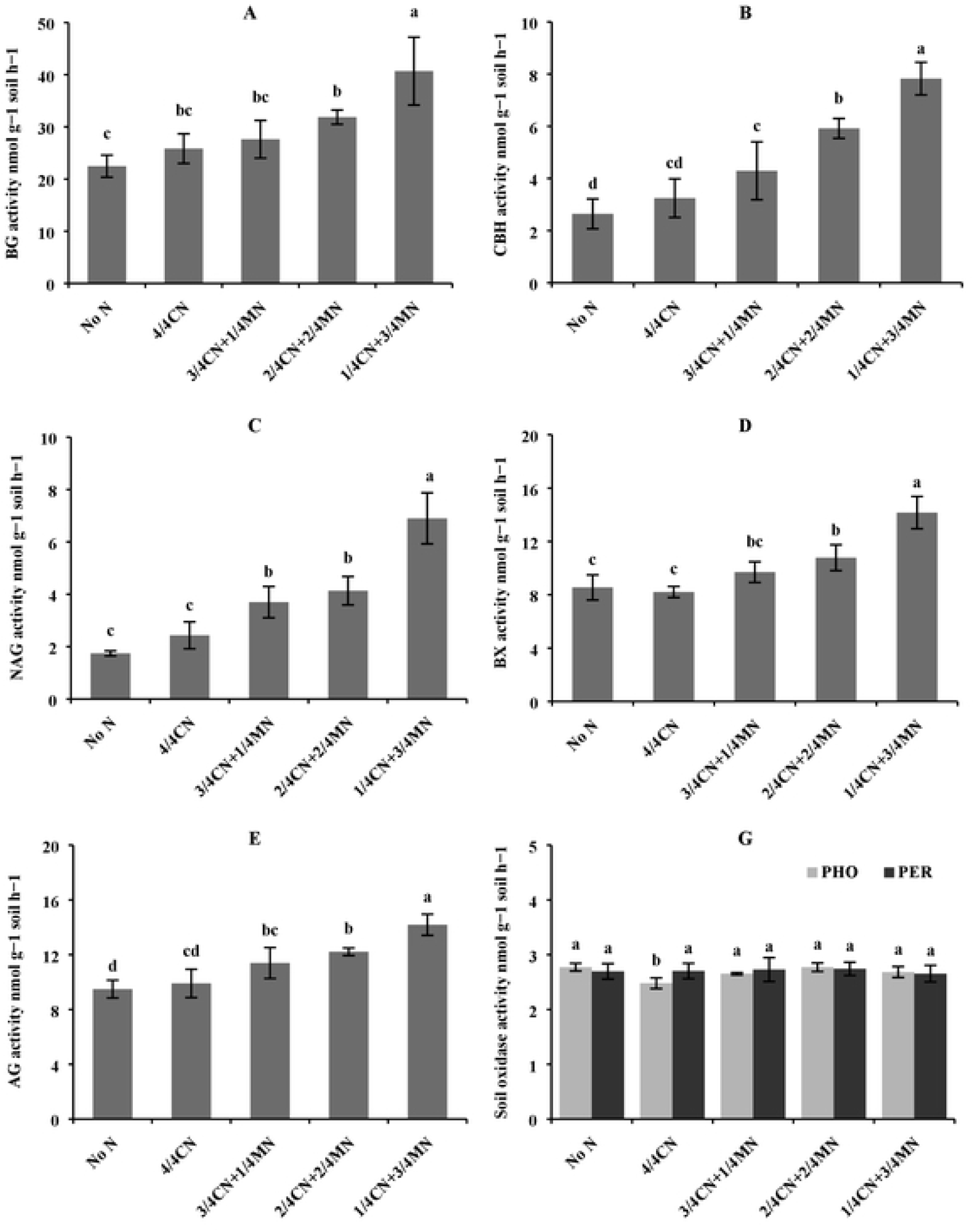
Changes in soil extracellular enzyme activities under different fertilization treatments. Vertical error bars represent the standard deviation (n = 3), and lowercase letters indicate significant differences among different treatments at the *p* < 0.05 level.

Compared with chemical fertilization treatments, the values of Gmea, GH and GH/GOR under manure-amended treatments were significantly increased by 17.81%–75.20%, 23.89%–121.32% and 19.22%–126.32%, respectively (Table 4). The values of Gmea, GH and GH/GOR increased as the manure application rate increased. Additionally, no significant differences were observed in the GOR values among different fertilization treatments (2.59–2.76).

Results from principal component analysis (PCA) indicated that soil enzyme activities changed under different fertilization treatments (Fig. 3). The first principal component (PC1) explained 96.9%, and the second (PC2) explained 2.0%, of the total variance (Fig. 3a). Soil enzyme activities profiles showed significant separation among the five-fertilization treatments. Along PC1, higher manure-amended treatments (2/4CN+2/4MN and 1/4CN+3/4MN) had higher scores than chemical fertilization treatments. Along PC2, soil enzyme activities profile scores did not differ among the five treatments. The PC loadings for soil enzyme activities (Fig. 3b) and PC scores indicated that higher manure-amended treatments enhanced soil hydrolase (BG, CBH, NAG, BX and AG) activities, and the values of Gmea, GH and GH/GOR.

**Fig. 3.**
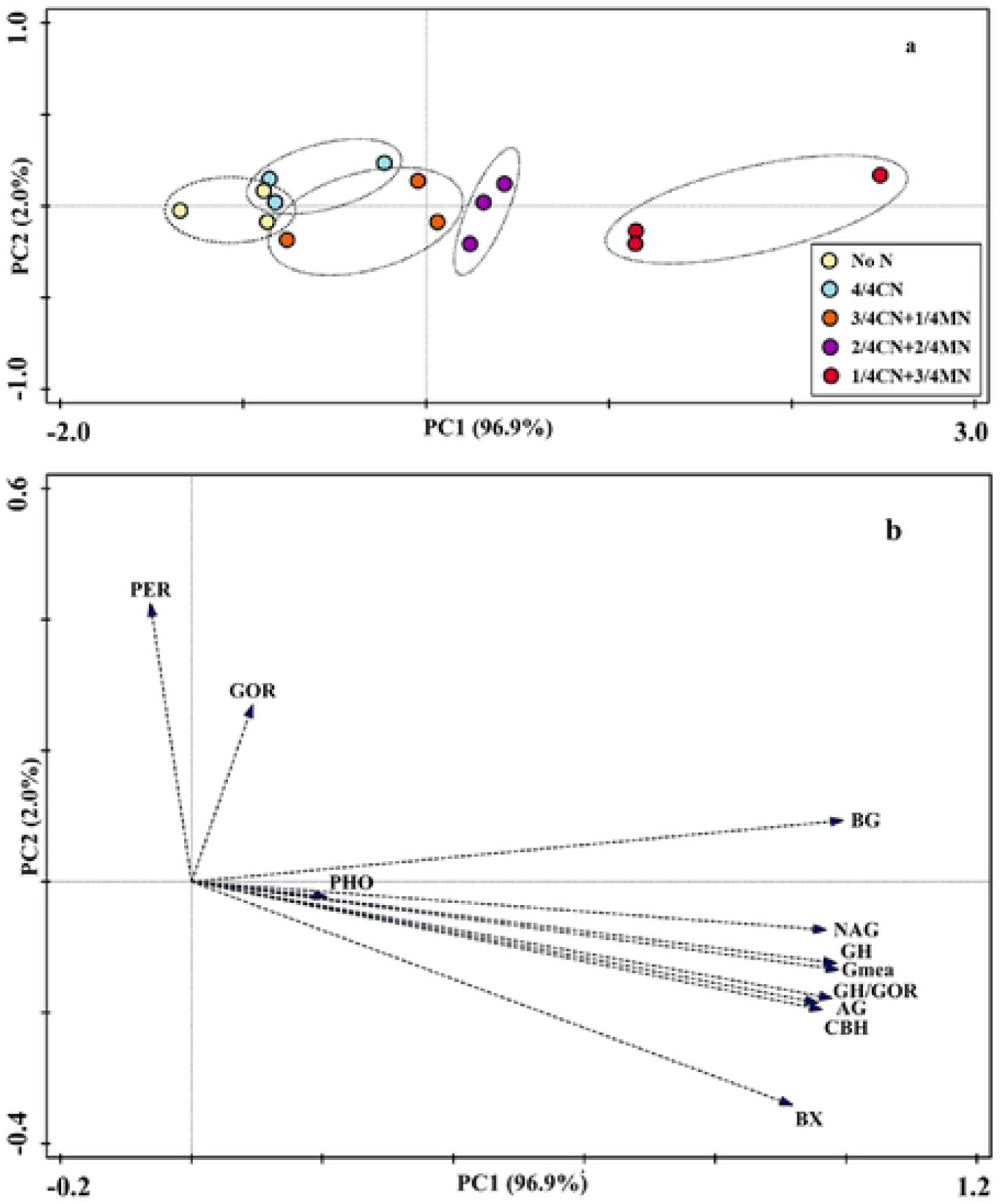
Principal component analysis (PCA) of soil extracellular enzyme activities profiles (a), and loading values for individual soil extracellular enzyme activities (b) from different fertilizer treatments.

### 3.3 Changes in soil microbial community structure

After celery harvest, total PLFA contents in manure-amended treatments (69.55–80.63 nmol g^−1^ soil) were significantly (*P* < 0.05) higher than those in chemical fertilization treatment (48.46–51.98 nmol g^−1^ soil), and total PLFA contents increased as the manure application rate increased (Fig. 4a). No significant differences were found in total PLFA contents between chemical fertilization treatments. Similar trends were observed for the contents of fungi (saprotrophic fungi (SF) and arbuscular mycorrhizal fungi (AMF)), bacteria (Gram-positive (G+) bacteria and Gram-negative (G−) bacteria) and actinomycete. Specific trends are as follows: 1/4CN+3/4MN > 2/4CN+2/4MN > 3/4CN+1/4MN > 4/4CN and No N (Fig. 4a, 4b, 4c and 4d).

**Fig. 4.**
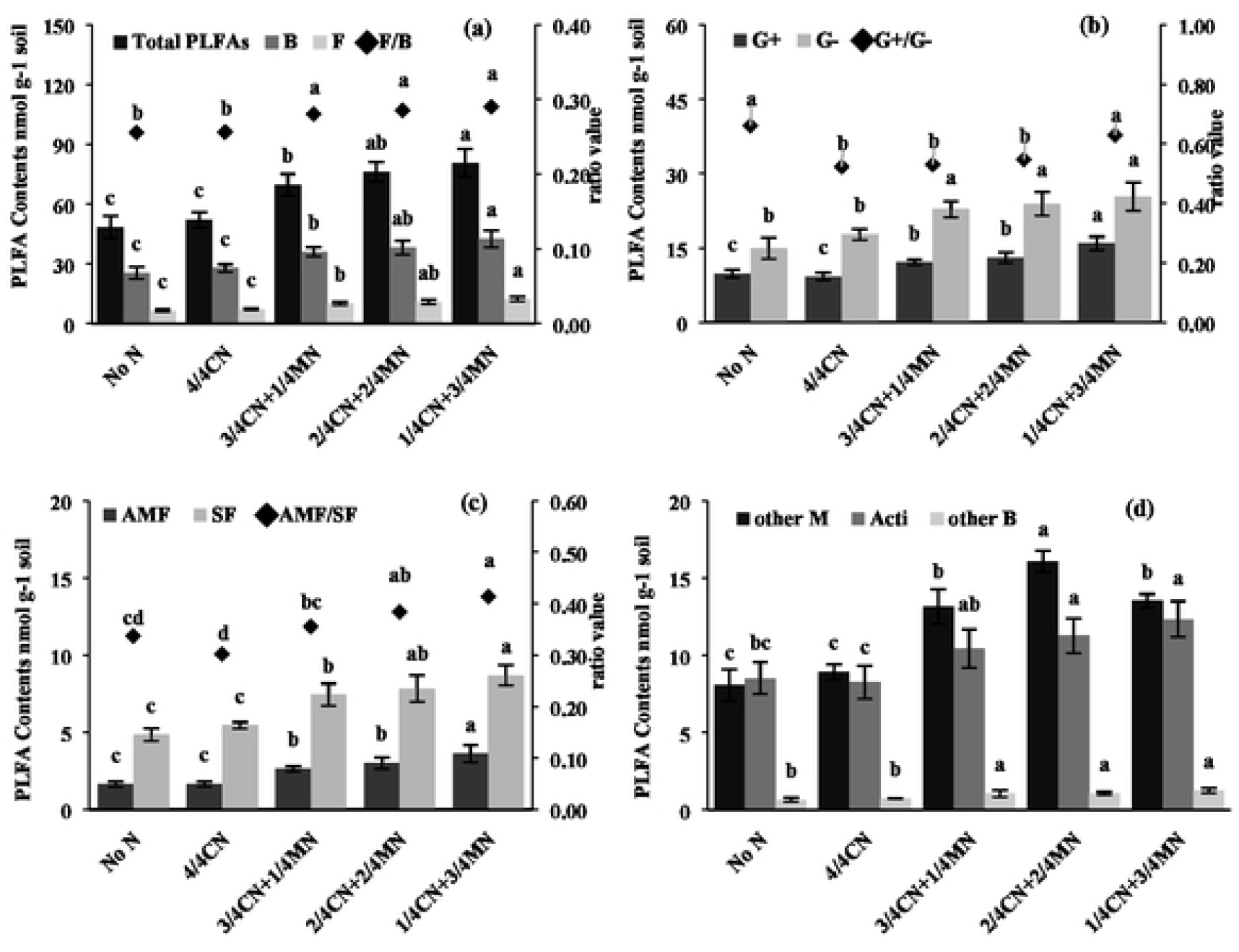
Changes in concentrations (nmol g^−1^ soil) of total PLFAs and microbial subgroups of PLFAs and associated ratios under different fertilization treatments. Vertical error bars represent the standard deviation (n = 3), and lowercase letters indicate significant differences among different treatments at the *p* < 0.05 level. Note: B bacteria, F fungi, G+: Gram-positive bacteria; G−: Gram-negative bacteria; AMF arbuscular mycorrhizal fungi, SF saprotrophic fungi, M microorganism, Acti actinomycetes.

The ratios of fungi to bacteria (F/B) and arbuscular mycorrhizal fungi to saprotrophic fungi (AMF/SF) in manure-amended treatments were significantly increased by 9.51%-13.48% and 5.55%-37.09%, respectively, compared to those in chemical fertilization treatments (Fig. 4a and 4c). Compared to 4/4CN treatment, the ratio of Gram-positive/Gram-negative bacteria (G+/G−) slightly increased under 3/4CN+1/4MN and 2/4CN+2/4MN, and significantly (*P* < 0.05) increased by 20.26% under 1/4CN+3/4MN. Additionally, the G+/G− values were highest in the No N treatment (0.66), and significantly higher by 26.33% than 4/4CN treatment (Fig. 4b).

The PCA of the 24 PLFAs data indicated that soil microbial community structure was markedly affected by manure application, but was not different (*P* > 0.05) between chemical fertilization treatments (No N and 4/4CN), indicated by their closest scores along the first principal component (PC1) and the second component (PC2) (Fig. 5a). The first two components, PC1 and PC2 explained 86.6% and 6.2% of the total variance in PLFAs profiles (Fig. 5). PC1 axis differentiated manure-amended treatments from chemical fertilization treatments, whereas PC2 axis did not differentiate different fertilization treatments. The PC loadings for individual PLFAs (Fig. 5b) and PC scores indicated that manure addition enhanced the vast majority of PLFAs contents.

**Fig. 5.**
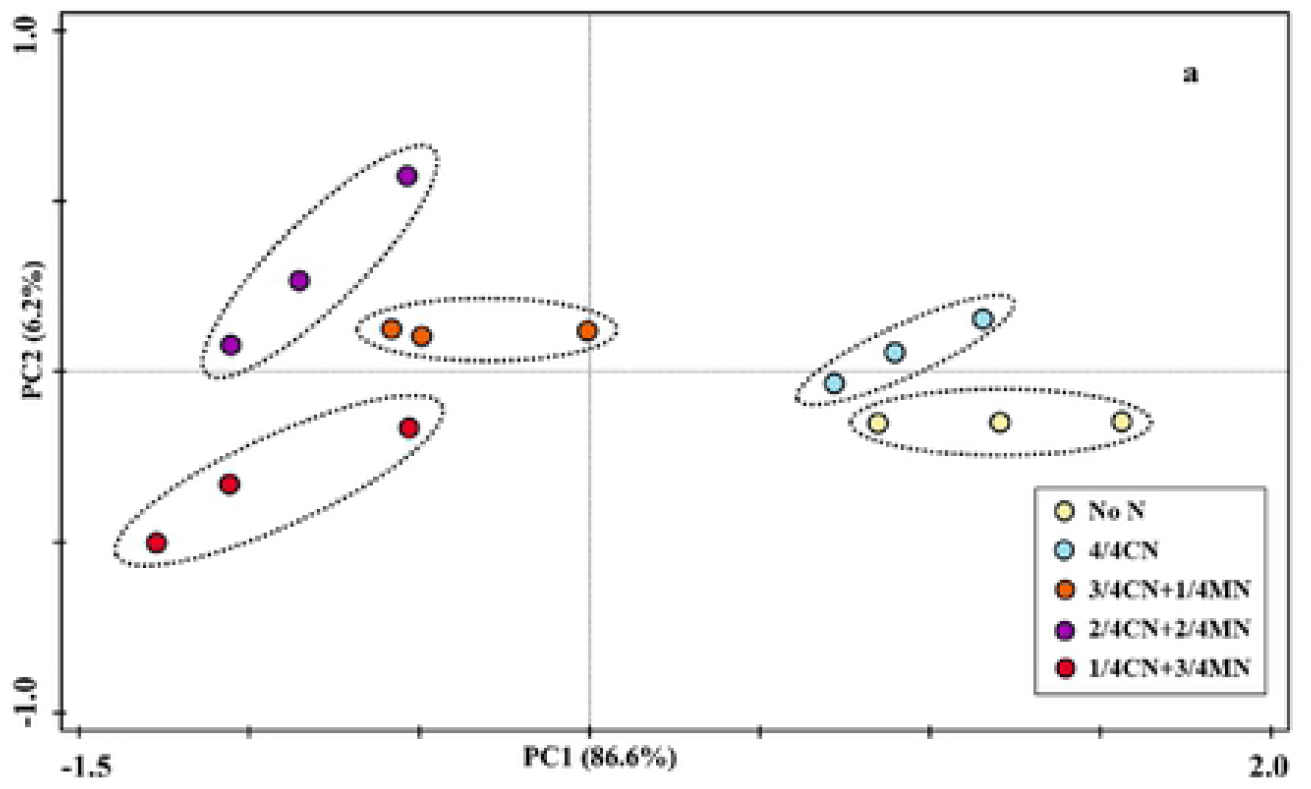
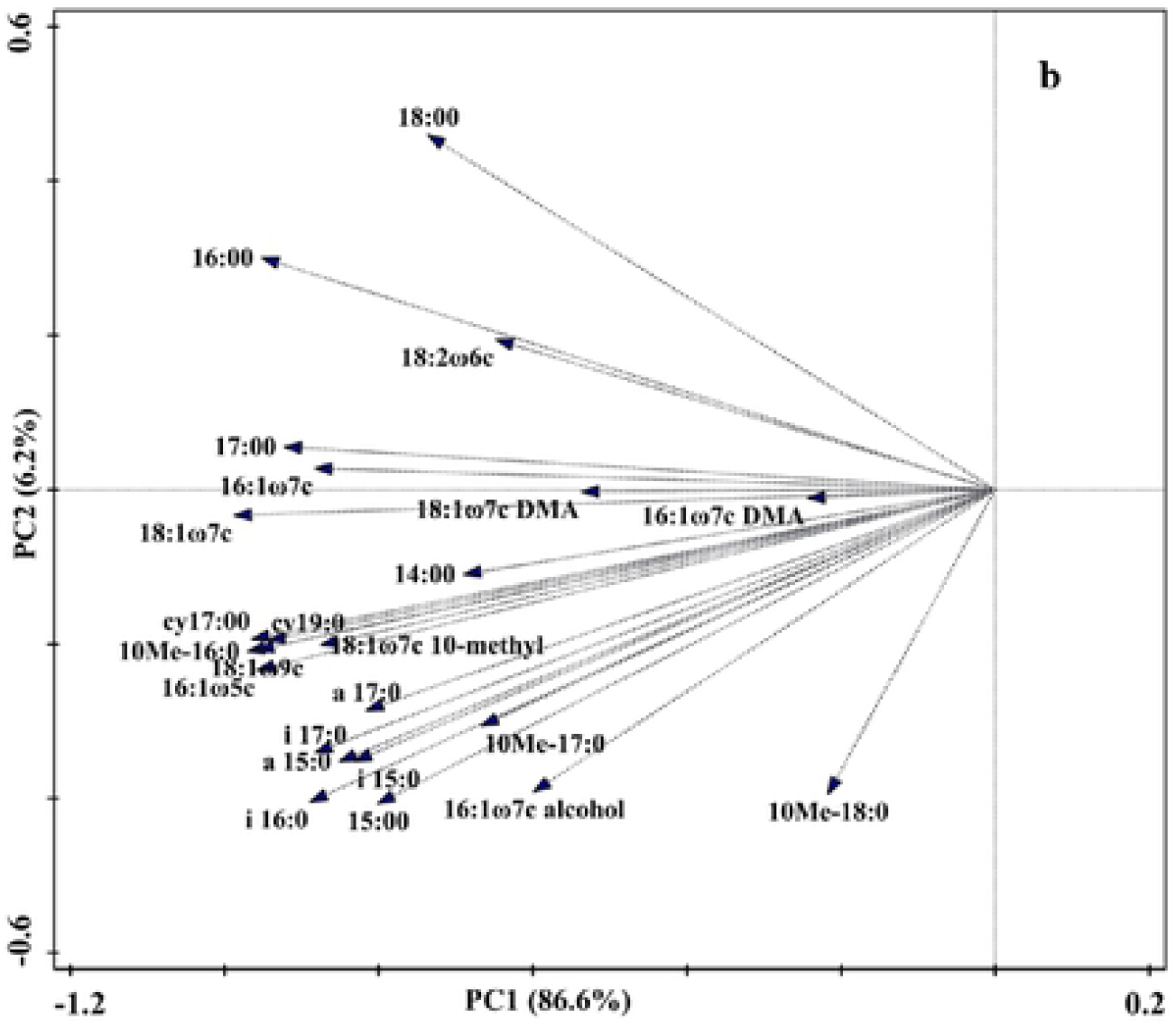
Principal component analysis (PCA) of soil phospholipid fatly acids (PLFAs) (a), and loading values for individual PLFAs (b) from different fertilizer treatments.

### 3.4 Changes in soil microbial community diversity and physiological stress indices

Soil microbial community and functional (enzyme) diversity in this study varied among different fertilization treatments (Table 5). Compared to chemical fertilization treatments, Shannon–Wiener diversity index (H’_M_) and Margalef richness index (SR) slightly increased under 3/4CN+1/MN treatment, significantly (*P* < 0.05) increased under 2/4CN+2/4MN and 1/4CN+3/MN treatment by 1.44%–3.74% and 10.42%–17.15%, respectively. Contrary to H’_M_ and SR, higher rates of manure input (1/4CN+3/4MN and 2/4CN+2/4MN) significantly reduced Pielou evenness index (J) by 2.58%–6.10% than chemical fertilization treatments. Soil enzyme function diversity index (H’_E_) calculated from all enzyme activities showed the similar tendency as the H’_M_: 1/4CN+3/4MN (1.94) > 2/4CN+2/4MN (1.93) > 3/4CN+1/4MN (1.91) > 4/4CN (1.88) > No N (1.84).

The cy17:0/16:1ω7c values were highest in the No N treatment (0.377), followed by manure-amended treatments (0.285–0.316), and the lowest value was observed in the 4/4CN treatment (0.223) (Fig. 6). Compared to the 4/4CN treatment, the (i17:0 + i15:0)/(a17:0 + a15:0) values slightly increased under 3/4CN+1/4MN treatment, significantly (*P* < 0.05) increased by 8.16% under 2/4CN+2/4MN treatment and by 13.89% under 1/4CN+3/4MN treatment.

**Fig. 6.**
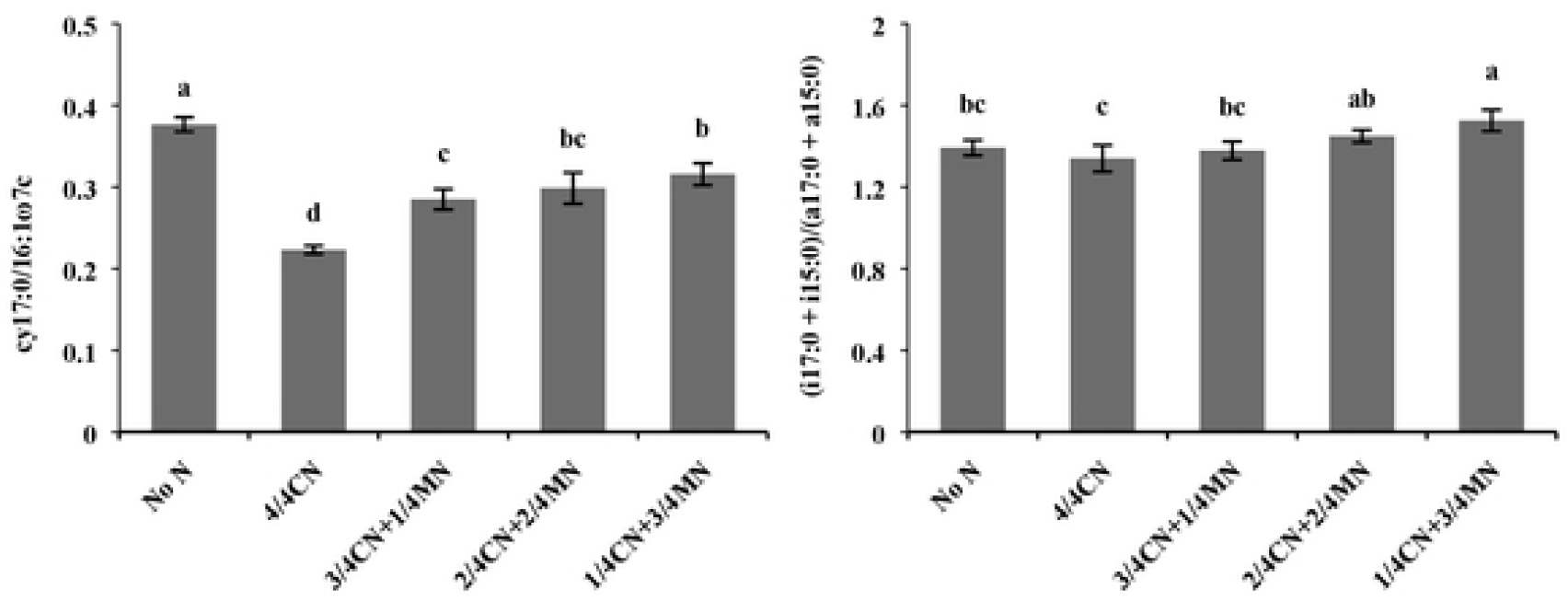
Changes in soil bacterial physiological stress indices under different fertilization treatments. Vertical error bars represent the standard deviation (n = 3), and lowercase letters indicate significant differences among different treatments at the *p* < 0.05 level.

### 3.5 Correlations between soil physicochemical properties and soil microbial characteristics

According to the Redundancy analysis (RDA) of the activities of all enzyme constrained by soil physicochemical properties, the first and second ordination RDA axis (RDA1 and RDA2) explained 87.8% and 5.6% of the total variance, respectively (Fig. 7a). The RDA confirmed that SOC content (F = 48.8, *P* = 0.002) and TN content (F = 3.0, *P* = 0.042) were strongly associated with soil enzyme activities and explained 78.9% and 4.2% of the total variance of enzyme activities data, respectively (Fig. 7a). Another RDA was performed using soil physicochemical properties as explanatory variables and soil PLFAs profiles as response variables (Fig. 7b). RDA1 and RDA2 accounted for 86.3% and 5.6% of the total variance of the PLFA data, respectively. Forward selection indicated that SOC content (F = 31.9, *P* = 0.002) was the most predominant variable, explaining 71.0% of the total variance of the PLFA data, followed by NH_4_^+^-N content (9.9%, F = 6.2, *P* = 0.012), AP content (5.9%, F = 4.9, *P* = 0.004) and soil EC (4.9%, F = 6.0, *P* = 0.006).

**Fig. 7.**
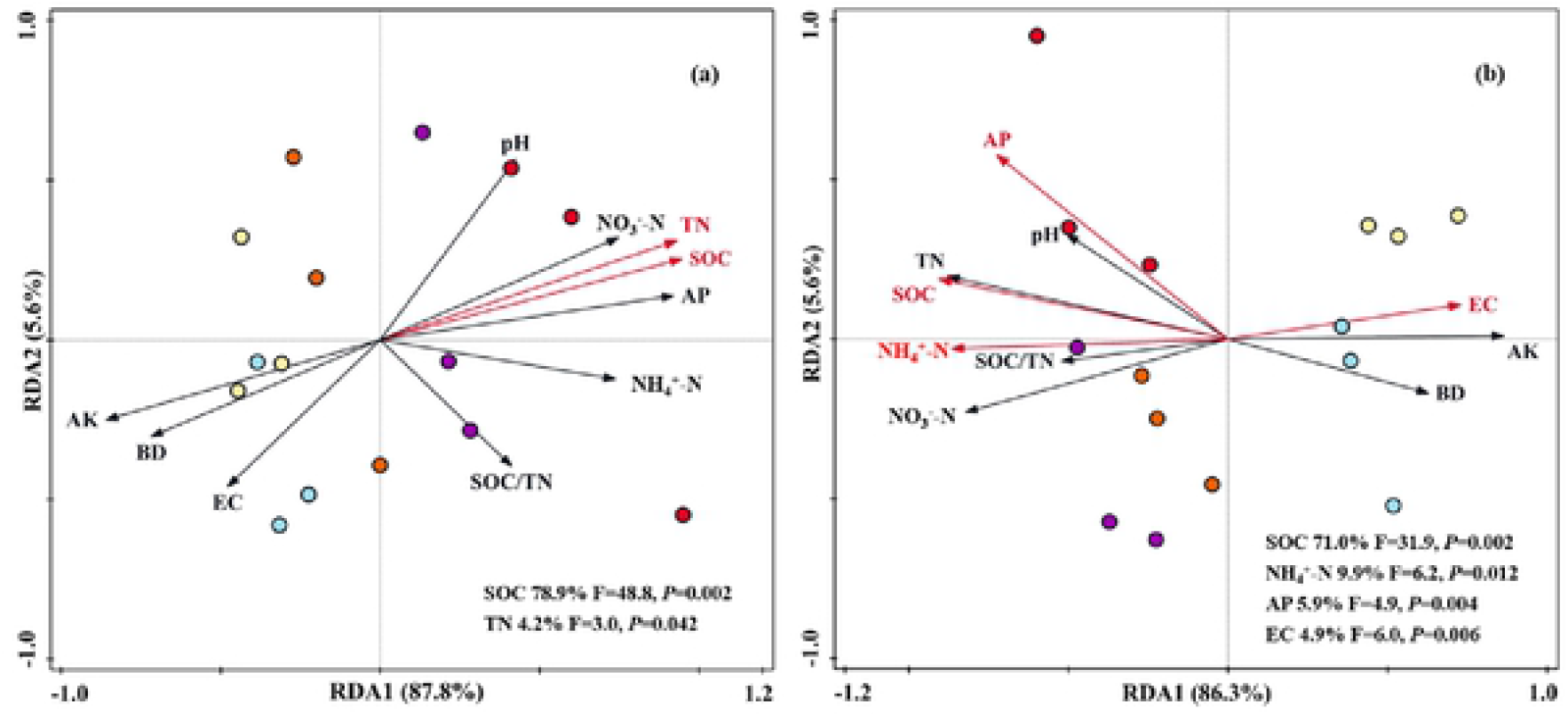
Redundancy analysis (RDA) of soil enzyme activities (a), microbial community structure (b) constrained by soil physicochemical properties under different fertilization treatments. Lengths and angles of the arrows indicate the correlations between selected soil physicochemical properties and soil enzyme activities, PLFA profiles. SOC soil organic caibon, TN total nitrogen, NO_3_^−^-N nitrate nitrogen, NH_4_^+^-N ammonium nitrogen, AP available phosphorus, AK available potassium and BD bulk density.

## 4. Discussion

### 4.1 Effects of manure substitution of chemical fertilizer on soil physicochemical properties

Organic amendments and substitution of chemical fertilizers are increasingly recommended as an important means to sustain crop yield and soil quality, which is attributed to changes in soil physicochemical properties and biological functions [21]. Our results indicated that manure addition improved soil nutrient-related properties, as evidenced by the increase in SOC, NO_3_^−^-N, NH_4_^+^-N and AP contents (Table 3), which was similar to previous studies [41]. These positive effects could be attributed to manure containing amounts of carbon resources, and subsequently release of these resources into soil [42]. Manure application could improve the nutrient retention ability of soil, which could explain the results described above [43]. Additionally, we found manure addition decreased soil AK contents. This is in contrast to most studies that annual addition of manure increased soil AK contents in double maize and maize–wheat double cropping systems [44, 45], but some researchers indicated that manure application markedly decreased soil AK contents in an intensive vegetable farm [46], which could be caused by massive consumption of K nutrients during vegetable growth [47]. Another possible explanation was that great demands for nutrients during microbial proliferation reduced short-term nutrients (K) availability [48]. In other words, manure application in this study promoted vegetable growth (Table S1) and soil microbial proliferation (Fig. 1 and Fig. 4), which consumed large amounts of nutrients (K) and therefore reduce soil nutrients (K) availability. Soil pH and EC values, as important indicators of soil chemical and electronical properties, can directly or indirectly affect the distribution of nutrient elements and the nutrient status of soil, thus playing vital roles on soil fertility and quality [16]. We found that manure application markedly increased soil pH values by 2.25%–2.77%, while markedly reduced soil EC values by16.67%–21.14%, compared to 4/4CN treatment. These results were supported by the findings of Wei et al. [49], who found that manure inputs alleviated the negative effect of chemical fertilizer application on soil pH. The possible explanation was that organic manure inputs could alleviate nitrification activities (acid-producing process) caused by chemical N inputs [50]. Mokolobate and Haynes [51] indicated organic manure contains humic-type substances with large amounts of carboxyl, phenolic, and enolic functional groups, which can consume the protons (e.g., Al^3+^, H^+^, etc.), this perhaps explained the reduce of soil EC values in manure-amended soils.

### 4.2 Effects of manure substitution of chemical fertilizer on soil enzyme activity

Soil extracellular enzyme activities (EEAs) are vital indicators of soil microbial activity, and they are closely associated with SOC decomposition and nutrient cycling [13]. On the basis of their functions, soil EEAs can be divided into hydrolases and oxidases that decompose substrates of various composition and complexity, which are mostly affected by fertilization [52]. It’s worth noting that the positive effects of manure application on soil hydrolase activities were supported by most scholars [17, 46], but the influences of manure application on soil oxidase activities were controversial [53, 54]. In our study, manure application was beneficial to enhance soil hydrolase (BG, CBH, NAG, BX and AG) activities (Fig. 2). The trends in soil hydrolase activities could be interpreted by soil characteristics related to carbon and nutrient availability, which are well known to be strongly influenced by fertilizer managements [22, 53].

Soil oxidases are generally produced by fungi, and their activities were different from soil hydrolase activities in soil [53, 55]. In our study, soil PHO and PER activities were not affected by manure application, whereas NPK chemical fertilizer application (4/4CN) markedly inhibited soil PHO activity relative to other treatments. Similar results have been reported in other studies [56, 57], they also found that chemical N inputs were shown to depress soil oxidase (PHO) activity. This could be because chemical N input could suppress the growth of fungi and therefore reduce the secretion of oxidases from fungi [58]. Finally, based on the data about geometric mean of enzyme activities, we summarized that manure application could significantly enhance soil hydrolase activities and had no effects on soil oxidase activities, thus driving shifts toward hydrolase-dominated (higher values of GH/GOR ratio) soil enzyme profile.

### 4.3 Effects of manure substitution of chemical fertilizer on soil microbial community composition

Soil microorganisms can be used as pivotal indicators of biogeochemical cycles because of their diversity, functional traits, dispersal ability and density [59]. PLFA analyses can provide information on several levels (“metrics”) of soil microbial community, from a whole-community profile to specific group abundance, microbial diversity, and physiological stress (Fig. 4, Fig. 6 and Table 5) [60]. Recently, lots of studies have investigated the response of soil community structure and abundance to manure application in different ecosystems. However, divergent and even conflicting results were often reported [23, 61]. In our study, all and individual microbial groups (e.g., fungi, bacteria and actinomycete, etc.) and microbial biomass (MBC and MBN) increased with increasing manure input rates, which was in accordance with the findings of Yue et al. [45]. This could be ascribed that higher manure inputs could provide greater diversity of available substrates for soil microorganism growth and reproduction [62]. Additionally, the microorganisms present in organic manure could also be contributing to the enhancement of soil microbial biomass [61].

Manure application not only increased all and individual microbial groups biomass, but also changed microbial community structure (Fig. 4). In line with the results of previous research on agricultural soil [63], we found that the F/B, G+/G− and AMF/SF ratios increased after 8 years manure inputs (Fig. 4). These changes indicated that some species (fungi, G+ bacteria and AMF) adapted better to manure application than other species (bacteria, G− bacteria and SF). On the basis of previous studies, the utilization of microorganisms for exogenous organic materials usually has a community succession effect: fast-growing G− bacteria proliferate soon after organic materials addition, and then it decreases, promoting the growth of other more slowly-growing microorganisms such as G+ bacteria or fungi [64]. Moreover, manure application could promote soil aggregation and create more big pores, which facilitate fungal growth [65]. This perhaps explained the increase of G+/G− and F/B ratios in manure-amended soils. It has been concluded in previous studies that different microorganisms have different effects on nutrient cycling and SOC accumulation in soil, and higher F/B, G+/G− and AMF/SF ratios are favorable for SOC accumulation and soil aggregation [66, 67]. Therefore, we speculated that manure application would optimize soil microbial community structure, and be beneficial for soil fertility and biological quality.

### 4.4 Effects of manure substitution of chemical fertilizer on soil microbial community diversity and physiological stress

Here, soil microbial community diversity and physiological stress were affected by manure application (Table 5 and Fig. 6). Ge et al. [68] and Yu et al. [69] found that the values of H’_M_ and SR were increased upon manure inputs in different agricultural soils of Northern China. Similarly, our results indicated that manure application not only increased soil microbial community diversity (H’_M_) and richness (SR), but also reduced soil microbial community evenness (Table 5). It has been suggested that manure application could (1) improve soil structure and (2) increase carbon and nutrient resource diversity, and subsequently improve soil microbial community diversity and richness [62, 70]. Interestingly, different microorganisms have been reported to be differently affected by fertilization, owing to their diverse structure and physiology (e.g., manure addition was more favorable to the growth of some species, such as fungi etc.) [71, 72]. This could be the reason that soil microbial community evenness was decreased in manure-amended soils.

The cy17:0/16:1ω7c and (i17:0 + i15:0)/(a17:0 + a15:0) ratios were denoted as bacterial physiological stress indices, which indicated microbial physiological status in response to environmental stress [72]. Higher values of these indices are associated with decreased bacterial growth rate and increased SOC and nutrient limitation [37]. In our study, the higher values of bacterial stress indices suggested that nutritional or environmental stress was more limiting to the growth of bacteria in the manure-amended soils. This is in contrast to usual expectation that organic fertilizers additions would decrease the values of these indices [72], but there are studies support our conclusions [73]. Our results could be caused by competition between fungi and bacteria for organic substrate and nutrients [74]. Namely, rapid and massive proliferation of fungi after the addition of manure had antagonistic effects on bacterial growth.

### 4.5 Linking soil microbial community composition and enzyme activity to soil physicochemical properties

Understanding the main environmental drivers regulating microbial communities and their functions represents an important challenge in microbial ecology [49]. RDA (Fig. 7a) indicated that the majority of the variation in soil enzyme activities could be explained by soil nutrient characteristics such as SOC contents (78.9%) and TN contents (4.2%). In agreement with our results, Wang et al. [32] found that the changes of soil enzyme activities were strongly related to soil total C and N in tropical forests. This reason maybe that sufficient C and N resources (derived from manure) could alleviate C and N limitation of soil microbial metabolism, and thus enhanced the secretion of enzymes [75]. It has been found in many studies that environmental factors and soil management practices largely determine soil microbial structure and composition [76]. The RDA results (Fig. 7b) demonstrated that SOC content (71.0%), NH_4_^+^-N content (9.9%), AP content (5.9%) and soil EC (4.9%) played important roles in shaping soil microbial community composition. This means that manure addition resulted in lower soil EC and improved soil chemical properties (SOC, NH_4_^+^-N and AP), thus induced a strong response of the changes of soil microbial structure. Luo et al. [63] and Ma et al. [77] also reported that C and nutrient (N and P) supplied by manure were major influence factors affecting soil microbial community structure in semiarid farmland and subtropical paddy soils, respectively. In conclusion, the changes in soil microbial characteristics were driven by different fertilization patterns induced changes in soil physicochemical properties: the ten selected environmental factors together accounted for >90% of the total variation (Fig. 7a and 7b), while SOC and nutrient (N) availability (especially SOC), rather than other soil physicochemical properties, are the vital drivers among them.

## 5. Conclusions

The 8-year field experiment clearly revealed that the responses of soil physicochemical properties, soil enzyme activities and microbial community structure to different fertilization patterns in a GVP of Tianjin, China. We confirmed that manure substitution of chemical fertilizer, especially manure application at higher rates, (1) improved soil physicochemical properties and potential crop productivity, (2) enhanced soil enzyme activities, promoted soil microbial growth, altered microbial community composition (increased the F/B, AMF/SF, and G+/G− ratios) and optimized soil microbial community structure, and (3) improved soil microbial community diversity and increased bacterial physiological stress. Meanwhile, we found that SOC content and nutrient (N)-related properties (especially SOC) played important roles in determining the effects of different fertilization patterns on the changes of soil microbial characteristics. In conclusion, the practices of organic manure substitution of chemical fertilizer, especially at high manure application rates (20.8–31.2 Mg ha^−1^ yr^−1^), could be a promising approach to develop sustainable agriculture, and was beneficial for vegetable development and soil quality in GVPs of China.

## Acknowledgements

This research was supported by the National Key Research and Development Program of China (2016YFD0201001), China Agriculture Research System (CARS-23-B02), and the Key Research and Development Program of Shandong Province, China (2017CXGC0206). We thank Gareth Thomas, PhD, from Liwen Bianji, Edanz Group China (www.liwenbianji.cn/ac), for editing the English text of a draft of this manuscript.

